# Outer membrane lipid homeostasis via retrograde phospholipid transport in *Escherichia coli*

**DOI:** 10.1101/109884

**Authors:** Rahul Shrivastava, Xiang’Er Jiang, Shu-Sin Chng

**Affiliations:** Department of Chemistry, National University of Singapore, Singapore 117543; Singapore Center for Environmental Life Sciences Engineering, National University of Singapore (SCELSE-NUS), Singapore 117456

**Keywords:** outer membrane stability, membrane homeostasis, lipid trafficking, membrane lipid asymmetry, membrane contact sites, TolQRA

## Abstract

Biogenesis of the outer membrane (OM) in Gram-negative bacteria, which is essential for viability, requires the coordinated transport and assembly of proteins and lipids, including lipopolysaccharides (LPS) and phospholipids (PLs), into the membrane. While pathways for LPS and OM protein assembly are well-studied, how PLs are transported to and from the OM is not clear. Mechanisms that ensure OM stability and homeostasis are also unknown. The trans-envelope Tol-Pal complex, whose physiological role has remained elusive, is important for OM stability. Here, we establish that the Tol-Pal complex is required for PL transport and OM lipid homeostasis in *Escherichia coli*. Cells lacking the complex exhibit defects in lipid asymmetry and accumulate excess phospholipids (PLs) in the OM. This imbalance in OM lipids is due to defective retrograde PL transport in the absence of a functional Tol-Pal complex. Thus, cells ensure the assembly of a stable OM by maintaining an excess flux of PLs to the OM only to return the surplus to the inner membrane. Our findings also provide insights into the mechanism by which the Tol-Pal complex may promote OM invagination during cell division.

## Introduction

Lipid bilayers define cellular compartments, and thus life itself, yet our understanding of the assembly and maintenance of these structures are limited. In Gram-negative bacteria, the outer membrane (OM) is essential for growth, and allows the formation of an oxidizing periplasmic compartment beyond the cytoplasmic or inner membrane (IM) (Nikaido, 2003). The OM is asymmetric, with lipopolysaccharides (LPS) and phospholipids (PLs) found in the outer and inner leaflets, respectively. This unique lipid asymmetry is required for the OM to function as an effective and selective permeability barrier against toxic substances, rendering Gram-negative bacteria intrinsically resistant to many antibiotics, and allowing survival under adverse conditions. The assembly pathways of various OM components, including LPS (Okuda *et al*., 2016), β-barrel OM proteins (OMPs) (Hagan *et al.*, 2011), and lipoproteins (Okuda and Tokuda, 2011), have been well-characterized; however, processes by which PLs are assembled into the OM have not been discovered. Even though they are the most basic building blocks of any lipid bilayer, little is known about how PLs are transported between the IM and the OM. Unlike other OM components, PL movement between the two membranes is bidirectional (Donohue-Rolfe and Schaechter, 1980; Jones and Osborn, 1977; Langley *et al*., 1982). While anterograde (IM-to-OM) transport is essential for OM biogenesis, the role for retrograde (OM-to-IM) PL transport is unclear. How assembly of the various OM components are coordinated to ensure homeostasis and stability of the OM is also unknown.

The Tol-Pal complex is a trans-envelope system highly conserved in Gram-negative bacteria (Lloubes *et al*., 2001; Sturgis, 2001). It comprises five proteins organized in two sub-complexes, TolQRA in the IM and TolB-Pal at the OM. In *Escherichia coli*, these sub-complexes interact in a proton motive force (pmf)-dependent fashion, with TolQR transducing energy to control conformational changes in TolA and allowing it to reach across the periplasm to contact Pal (Cascales *et al*., 2000; Germon *et al*., 2001), an OM lipoprotein that binds peptidoglycan (Godlewska *et al*., 2009). TolA also interacts with periplasmic TolB (Walburger *et al*., 2002), whose function within the complex is not clear. The TolQRA sub-complex is analogous to the ExbBD-TonB system (Lloubes *et al*., 2001; Cascales *et al*., 2001; Witty *et al*., 2002), where energy-dependent conformational changes in TonB are exploited for the transport of metal-siderophores across the OM (Gresock *et al*., 2015). Unlike the ExbBD-TonB system, however, the physiological role of the Tol-Pal complex has not been elucidated, despite being discovered over four decades ago (Bernstein *et al*., 1972; Lazzaroni and Portalier, 1981). The Tol-Pal complex has been shown to be important for OM invagination during cell division (Gerding *et al*., 2007), but mutations in the *tol-pal* genes also result in a variety of phenotypes, such as hypersensitivity to detergents and antibiotics, leakage of periplasmic proteins, and prolific shedding of OM vesicles, all indicative of an unstable OM (Lloubes *et al*., 2001). In addition, removing the *tol-pal* genes causes envelope stress and up-regulation of the σ^E^ and Rcs phosphorelay responses (Vines *et al*., 2005; Clavel *et al*., 1996). It has thus been suggested that the Tol-Pal complex may in fact be important for OM stability and biogenesis. Interestingly, the *tol-pal* genes are often found in the same operon as *ybgC* (Sturgis, 2001), which encodes an acyl thioesterase shown to interact with PL biosynthetic enzymes in *E. coli* (Gully and Bouveret, 2006). This association suggests that the Tol-Pal complex may play a role in PL metabolism and/or transport.

Here, we report that the Tol-Pal complex is required for retrograde PL transport and OM lipid homeostasis in *E. coli*. We show that cells lacking the Tol-Pal complex exhibit defects in OM lipid asymmetry, as judged by the presence of outer leaflet PLs. We further demonstrate that *tol-pal* mutants accumulate excess PLs (relative to LPS) in the OM, indicating lipid imbalance in the membrane. Finally, using OM PL turnover as readout, we establish that the Tol-Pal complex is functionally important for efficient transport of PLs from the OM back to the IM. Our work solves a longstanding question on the physiological role of the Tol-Pal complex, and provides novel mechanistic insights into lipid homeostasis in the OM.

## Results

### Cells lacking the Tol-Pal complex exhibit defects in OM lipid asymmetry

To elucidate the function of the Tol-Pal complex, we set out to characterize the molecular nature of OM defects observed in *tol-pal* mutants in *E. coli*. Defects in the assembly of OM components typically lead to perturbations in OM lipid asymmetry (Wu *et al*., 2006; Ruiz *et al*., 2008). This is characterized by the accumulation of PLs in the outer leaflet of the OM, which serve as substrates for PagP-mediated acylation of LPS (lipid A) (Bishop, 2005). To determine if *tol-pal* mutants exhibit defects in OM lipid asymmetry, we analyzed lipid A acylation in strains lacking any member of the Tol-Pal complex. We demonstrated that each of the mutants accumulate more hepta-acylated lipid A in the OM compared to wild-type (WT) cells (Fig. 1). This OM defect, and the resulting SDS/EDTA sensitivity in these *tol-pal* mutants, are all corrected in the complemented strains (Fig. S1). We also examined other strains with known OM permeability defects. We detected increased lipid A acylation in strains with either impaired OMP (*bamB*, *bamD*, Δ*surA*) or LPS (*lptD4213*) biogenesis, as would be expected, but not in strains lacking covalent tethering between the cell wall and the OM (Δ*lpp*) (Fig. 1). Even though the Δ*lpp* mutant is known to exhibit pleiotropic phenotypes (Yem and Wu, 1978; Bernadac *et al*., 1998), it does not have perturbations in OM lipid asymmetry. In contrast to OMP or LPS assembly mutants, *tol-pal* strains produce WT levels of major OMPs and LPS in the OM (Fig. S2). These results indicate that *tol-pal* mutations lead to accumulation of PLs in the outer leaflet of the OM independent of OMP and LPS biogenesis pathways.

**Fig. 1.**
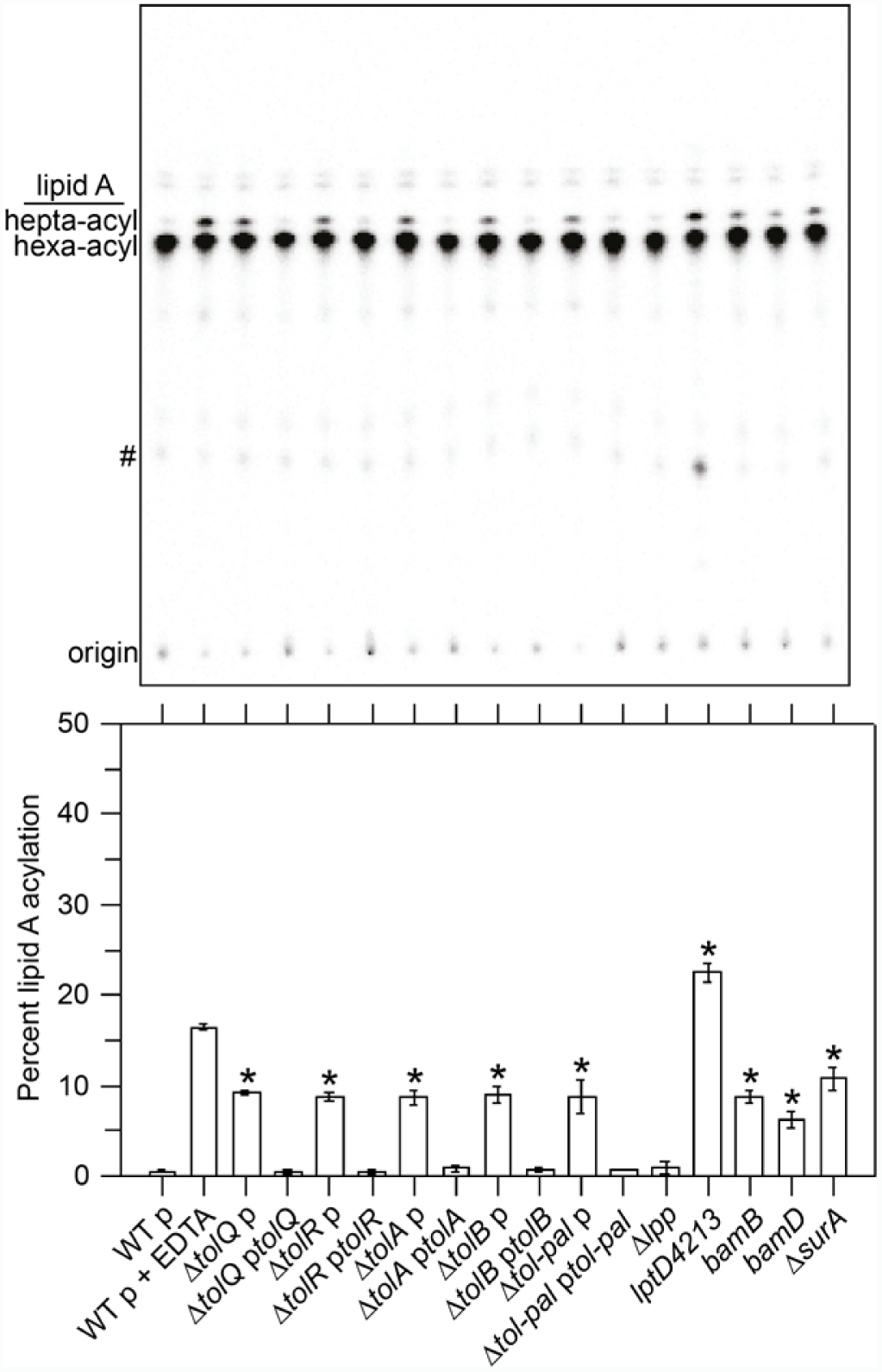
Cells lacking the Tol-Pal complex accumulate PLs in the outer leaflet of the OM as judged by lipid A acylation. Thin layer chromatographic (TLC) analysis of [^32^P]-labelled lipid A extracted from WT, Δ*tol pal*, and various mutant strains (*see text*). Where indicated, WT and *tol-pal* mutants contain an empty pET23/42 plasmid (p) (Wu *et al*., 2006) or one expressing the corresponding *tol-pal* gene(s) at low levels (e.g. p*tol-pal*). As a positive control for lipid A acylation, WT cells were treated with EDTA (to chelate Mg^2+^ and destabilize the LPS layer) prior to extraction. Equal amounts of radioactivity were spotted for each sample. Lipid spots annotated # represent 1 pyrophosphoryl-lipid A. Average percentages of lipid A acylation and standard deviations were quantified from triplicate experiments and plotted below. Student’s t-tests: * *p* < 0.005 as compared to WT.

### Cells lacking the Tol-Pal complex have disrupted OM lipid homeostasis

We hypothesized that the loss of OM lipid asymmetry in *tol-pal* mutants is due to defects in PL transport across the cell envelope. To test this, we examined the steady-state distribution of PLs (specifically labelled with [^3^H]-glycerol) between the IM and the OM in WT and *tol-pal* strains. We established that *tol-pal* mutants have ∼1.4-1.6-fold more PLs in their OMs (relative to the IMs) than the WT strain (Fig. 2A and Fig. S4). To ascertain if this altered distribution of PLs between the two membranes was due to the accumulation of more PLs in the OMs of *tol-pal* mutants, we quantified the ratios of PLs to LPS (both lipids now labelled with [^14^C]-acetate) following OM isolation and differential extraction. *tol-pal* mutants contain ∼1.5-2.5-fold more PLs (relative to LPS) in their OMs, when compared to the WT strain (Fig. 2B and Fig. S5). Since *tol-pal* mutants produce WT LPS levels (Fig. S2), we conclude that strains lacking the Tol-Pal complex accumulate excess PLs in their OMs, a phenotype that can be corrected via genetic complementation (Fig. 2). Consistent with this idea, *tol-pal* mutants, unlike WT (Fuhrer *et al*., 2006), are able to survive the toxic effects of LPS overproduction (Fig. S6), possibly due to a more optimal balance of PLs to LPS in their OMs. Importantly, having excess PLs makes the OM unstable, which can account for lipid asymmetry defects (Fig. 1), and increased permeability of the OM in *tol-pal* mutants (Lloubes *et al*., 2001). It also explains why these strains produce more OM vesicles (∼34-fold higher than WT cells, albeit only at ∼5% of total membranes (Fig. S7A)) (Bernadac *et al*., 1998). Consistent with this idea, OM vesicles isolated from the Δ*tolA* mutant similarly contain an elevated ratio of PLs to LPS, when compared to that in the WT OM (Fig. S7B). Furthermore, cells lacking the Tol-Pal complex are on average shorter and wider than WT cells (when grown under conditions with no apparent division defects) (Gerding *et al*., 2007); this reflects an increase in surface area of the rod-shaped cells, perhaps a result of increase in OM lipid content. As expected, we did not observe disruption of lipid homeostasis in the Δ*lpp* mutant (Fig. 2). However, we observed higher PL content in the OMs of strains defective in OMP assembly. We reasoned that this increase may help to stabilize the OM by filling the voids created by the decrease in properly-assembled OMPs. Since strains lacking the Tol-Pal complex have proper OMP assembly (Fig. 2A), the phenotype of excess PL build-up in the OM must be due to a different problem. Our results suggest that *tol-pal* mutations directly affect PL transport processes, and therefore OM lipid homeostasis.

**Fig. 2.**
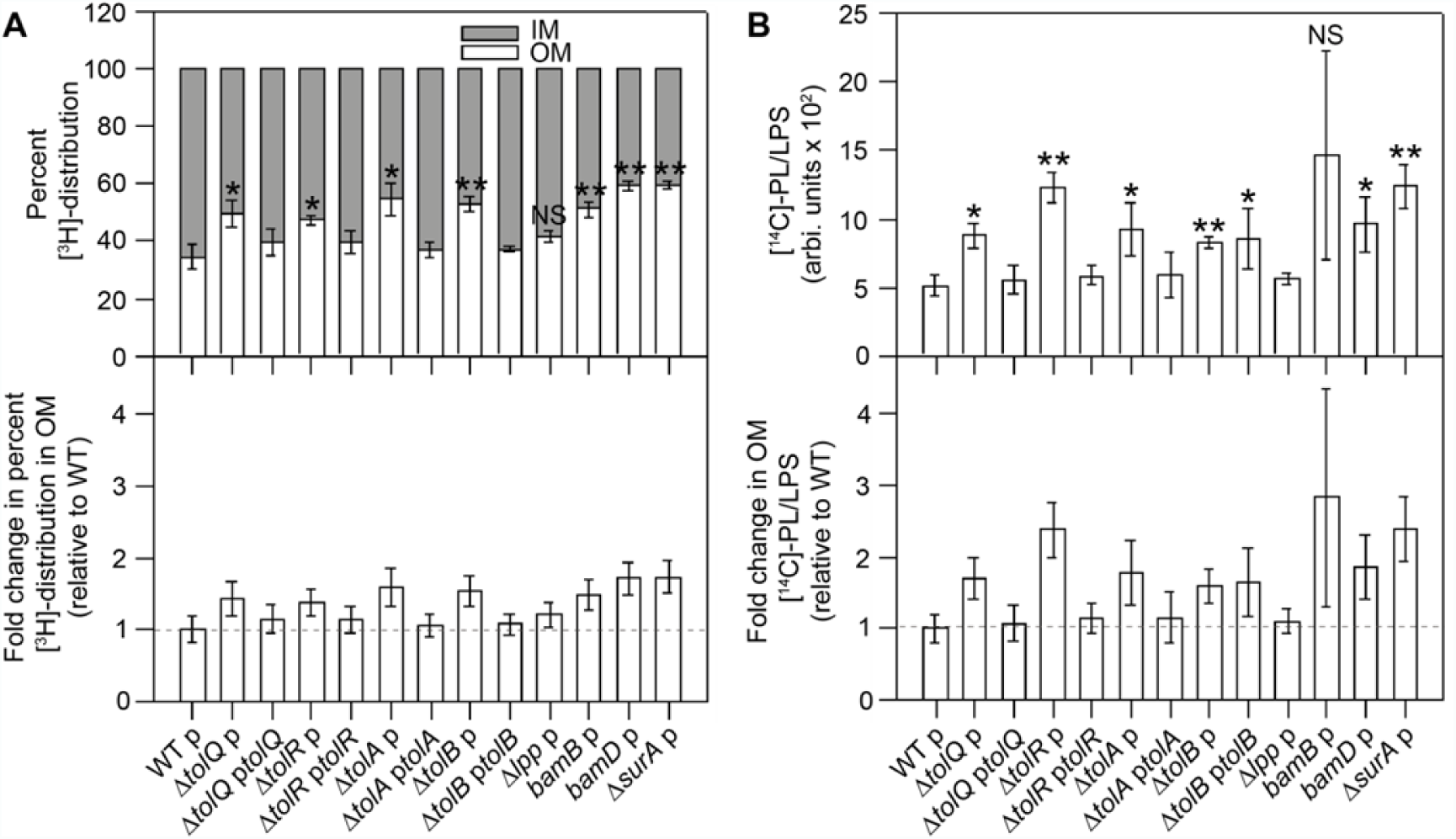
Cells lacking the Tol-Pal complex accumulate excess PLs (relative to LPS) in the OM. A. Steady-state distribution of [^3^H]-glycerol labelled PLs between the IM and the OM of WT, Δ*tol-pal*, and various mutant strains (*upper panel*)(Fig. S4). Distribution of [^3^H]-labelled PLs in the OMs of respective mutants expressed as fold changes relative to the WT OM (*lower panel*). The IMs and OMs from both WT and *tol-pal* mutants were separated with equal efficiencies during sucrose density gradient fractionation (Fig. S3). B. Steady-state PL:LPS ratios in the OMs of WT, Δ*tol-pal*, and various mutant strains (*upper panel*). Lipids were labelled with [^14^C]-acetate and differentially extracted from OMs (Fig. S5). OM PL:LPS ratios of respective mutants expressed as fold changes relative to that in the WT OM (*lower panel*). Error bars represent standard deviations calculated from triplicate experiments. Student’s t-tests: * *p* < 0.05; ** *p* < 0.005; NS, not significant (as compared to WT).

### Cells lacking the Tol-Pal complex are defective in retrograde PL transport

Unlike for other OM components, PL transport between the IM and the OM is bidirectional (Donohue-Rolfe and Schaechter, 1980; Jones and Osborn, 1977; Langley *et al*., 1982). Therefore, a simple explanation for the accumulation of excess PLs in the OMs of cells lacking the Tol-Pal complex is that there are defects in retrograde PL transport. To evaluate this possibility, we used the turnover of OM PLs (specifically anionic lipids, including phosphatidylserine (PS), phosphatidylglycerol (PG), and cardiolipin (CL)) as readout for the transport of PLs back to the IM (Fig. 3A). As an intermediate during the biosynthesis of the major lipid phosphatidylethanolamine (PE), PS is converted to PE by the PS decarboxylase (PSD) at the IM, and typically exists only at trace levels (Cronan, 2003). PG and CL have relatively short lifetimes (Kanfer and Kennedy, 1963; Kanemasa *et al*., 1967). While CL turnover is not well understood, PG turnover can occur via multiple pathways in *E. coli* (Hirschberg and Kennedy, 1972; Schulman and Kennedy, 1977; Yokoto and Kito, 1982). One specific way PG can turn over is by conversion to PE via PS, particularly when it is accumulated to abnormal levels in cells (Yokoto and Kito, 1982). Since all enzymatic activities possibly involved in converting PG to PS, and then to PE, are localized in the IM (Cronan, 2003), the turnover of OM anionic lipids via this pathway require, and therefore report on, retrograde PL transport (Fig. 3A). Such an assay has previously been employed to demonstrate retrograde transport for PS (Langley *et al*., 1982).

**Fig. 3.**
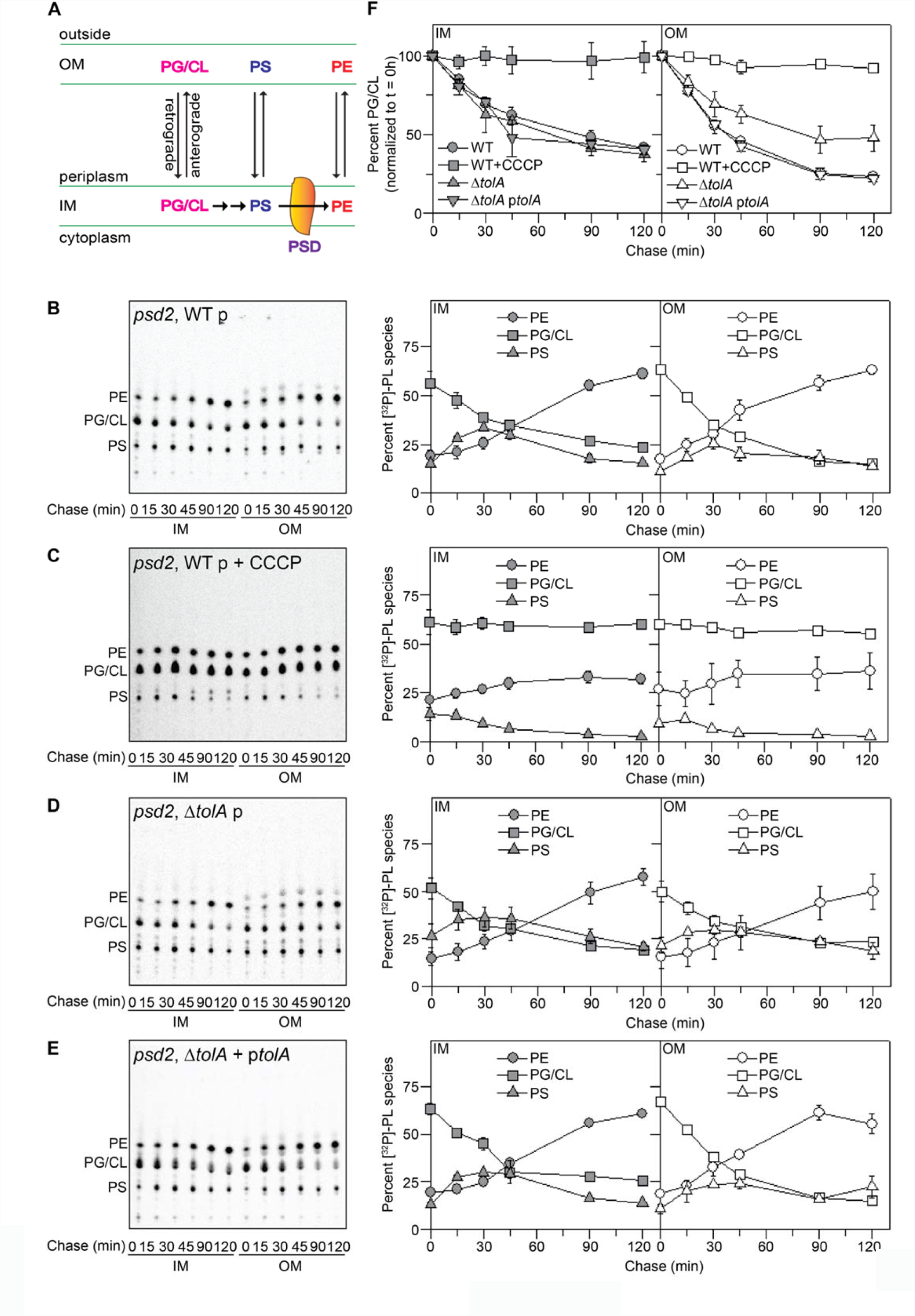
Cells lacking the Tol-Pal complex are defective in OM PG/CL turnover. A. A schematic diagram depicting bidirectional movement of PLs across the cell envelope, and the conversion of PG/CL to PE, via PS, in the IM. How PG may be converted to PS is not known, though one possible route may involve combining two PG molecules to give CL, and subsequent hydrolysis of CL to PG and PA (Audet, *et al.*, 1975), a precursor to all PLs (including PS) in cells. For clarity, other PG turnover pathways are also not shown. B-E. TLC time-course analyses of [^32^P]-pulse-labelled PLs extracted from the IMs and OMs of (B) WT, (C) WT (with CCCP added), (D) Δ*tolA*, and (E) *tolA*-complemented strains also harboring the *psd2* mutation. The average percentage levels of PE, PG/CL, and PS in the IM and OM at each time point, together with standard deviations, were quantified from triplicate experiments and shown on the right. F. The percentage levels of PG/CL in the IMs and OMs from (B-E) normalized to the corresponding levels at the start of the chase (0 min).

Using a strain expressing a temperature-sensitive (Ts) allele (*psd2*) of the gene encoding PSD (Hawrot and Kennedy, 1978), we pulse-labelled PLs with [^32^P]-phosphate at the restrictive temperature (42°C), and monitored the turnover of individual PL species in the OM during a chase period at the permissive temperature (30°C). At 42°C, the *psd2* strain accumulates substantial amounts of PS in both the IM and the OM (Fig. 3B, 0-min time point), as previously reported (Hawrot and Kennedy, 1978). With the restoration of PSD activity at 30°C, we observed initial increase but eventual conversion of PS to PE in both membranes (Fig. 3B, after 45-min time point), indicating that OM PS is transported back to the IM, converted to PE, and subsequently re-equilibrated to the OM (Langley *et al*., 1982). We also detected abnormally high PG/CL content in the *psd2* strain at 42°C, and saw rapid conversion of these lipids to PE in both membranes at 30°C (Fig. 3B), at rates comparable to what was previously reported (for PG) (Yokoto and Kito, 1982). The fact that PS levels increase initially but decrease after 45 min into the chase is consistent with the idea that PS is an intermediate along the turnover pathway for PG (Yokoto and Kito, 1982), as well as for CL. To confirm this observation, we also performed the chase at 42°C in the presence of a known PSD inhibitor (Satre and Kennedy, 1978) (these conditions completely shut down PSD activity), and found quantitative conversion of PG/CL to PS in both membranes (Fig. S8). We further showed that PG/CL-to-PE conversion is abolished in the presence of the pmf uncoupler carbonyl cyanide *m*-chlorophenyl hydrazone (CCCP) (Fig. 3C), demonstrating that cellular energy sources are required for this process (Yokoto and Kito, 1982), and that conversion occurs in the IM. The observation of PG/CL turnover in the IM is thus expected. The fact that we also observed the conversion of OM PG/CL to PE points towards an intact retrograde PL transport pathway for these lipids in the otherwise WT cells. Notably, turnover of OM PG/CL appears to be slightly faster than that of IM PG/CL (Fig. 3B), suggesting that retrograde transport of these lipids may be coupled to the turnover process.

We performed the same pulse-chase experiments with *psd2* cells lacking TolA. We detected PG/CL-to-PE conversion in the IM at rates comparable to WT (Fig. 3D, F; ∼67% and ∼71% PG/CL turnover at 2 h-chase in Δ*tolA* and WT IMs, respectively (Fig. 4A)), demonstrating that there are functional PG/CL turnover pathways in the Δ*tolA* mutant. In contrast, we observed substantial reduction of the turnover of OM PG/CL in these cells (Fig. 3D, F; ∼53% PG/CL turnover at 2 h-chase in the Δ*tolA* OM, compared to ∼79% for WT (Fig. 4A)), even though PS conversion to PE appears intact. These results indicate an apparent defect in the movement of PG and CL (but not PS) from the OM back to the IM, which is restored when complemented with functional *tolA*_*WT*_ (Fig. 3E, F, and Fig. 4A). Δ*tolR* mutant cells exhibit the same defect, and can similarly be rescued by complementation with functional *tolR*_*WT*_ (Fig. 4A). In contrast, no rescue was observed when Δ*tolR* was complemented using a *tolR* allele with impaired ability to utilize the pmf (*tolR*_*D23R*_) (Cascales *et al*., 2001) (Fig. 4A and Fig. S1); this indicates that Tol-Pal function is required for efficient PG/CL transport. We also examined PG/CL turnover in *psd2* cells lacking BamB, which accumulate excess PLs in the OM due to defects in OMP assembly (Fig. 2). Neither IM nor OM PG/CL turnover is affected (Fig. 4A), highlighting the different basis for OM PL accumulation in this strain compared to the *tol-pal* mutants. Our assay does not report on the retrograde transport of major lipid PE, which is relatively stable (Kanfer and Kennedy, 1963). However, since *tol-pal* mutants accumulate ∼1.5-fold more PLs in the OM (Fig. 2) without gross changes in PL composition (compared to WT) (Fig. S10), PE transport must also have been affected. We conclude that the Tol-Pal complex is required for the retrograde transport of bulk PLs in *E. coli*.

**Fig. 4.**
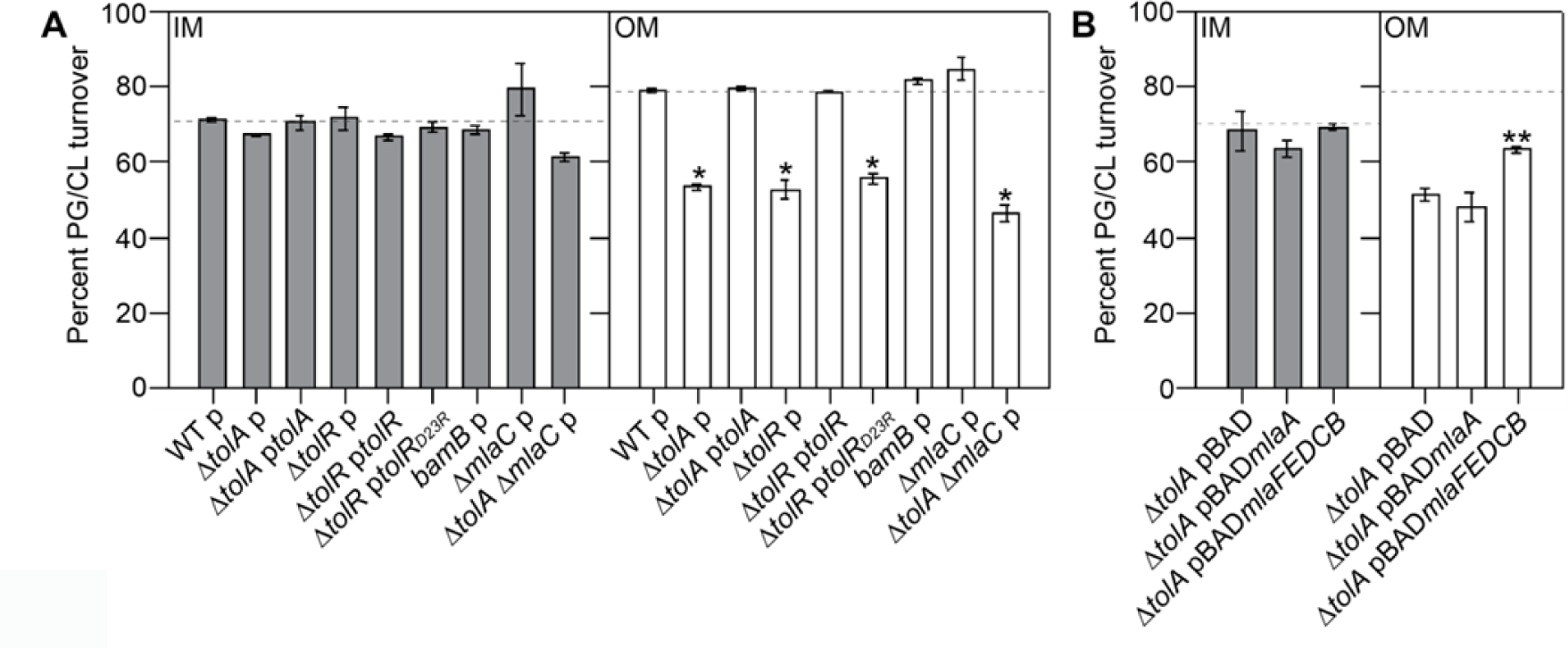
Tol-Pal function is required for efficient retrograde PG/CL transport, as judged by OM PG/CL turnover rates. Single time-point (2-h chase) quantification of the turnover rate of [^32^P]-labelled PG/CL in the IMs and OMs of (A) WT, *tol*-*pal* and various mutant strains, and (B) Δ*tolA* overexpressing OmpC-Mla components, all in the *psd2* background (*see text*) (Fig. S9). Percentage PG/CL turnover at 2-h is expressed as [(%PG/CL)_start_ – (%PG/CL)_2h_]/[(%PG/CL)_start_]. Average percentage lipid levels and standard deviations were quantified from triplicate experiments. Student’s t-tests: * *p* < 0.0005 as compared to WT; ** *p* < 0.0005 as compared to Δ*tolA*.

### Overexpressing a putative PL transport system partially rescues defects in retrograde PL transport observed in *tol-pal* mutants

Removing the Tol-Pal complex does not completely abolish retrograde PG/CL transport, indicating that there are other systems involved in this process. The OmpC-Mla system is important for the maintenance of OM lipid asymmetry, and is proposed to do so via retrograde PL transport (Malinverni and Silhavy, 2009; Chong *et al*., 2015). Two other related systems, the Pqi and Yeb systems, have recently been suggested to be involved in PL transport (Ekiert *et al.*, 2017); however, cells lacking either/both of these systems do not exhibit obvious OM defects unless the OmpC-Mla system is also removed (Nakayama and Zhang-Akiyama, 2016). To determine if the OmpC-Mla system plays a major role in retrograde PL transport in cells lacking the Tol-Pal complex, we examined OM PG/CL turnover in Δ*tolA* cells also lacking MlaC, the putative periplasmic lipid chaperone of the system. We first showed that cells lacking MlaC alone do not exhibit defects in OM PG/CL turnover (Fig. 4A). Evidently, removing MlaC also does not exacerbate the defects in retrograde PL transport in cells lacking the Tol-Pal complex, given that overall turnover rates of IM and OM PG/CL are similarly reduced in the double mutant. These results indicate that the OmpC-Mla system does not contribute significantly to retrograde transport of bulk lipids when expressed at physiological levels, as has been previously suggested (Malinverni and Silhavy, 2009). We also tested whether overexpressing the OmpC-Mla system can restore retrograde PL transport in *tol-pal* mutants. Interestingly, overexpression of MlaC and the IM MlaFEDB complex (Thong *et al*., 2016), but not MlaA, partially rescues OM PG/CL turnover in the Δ*tolA* mutant (Fig. 4B). However, this has no consequential effect on alleviating permeability defects observed in the Δ*tolA* strain (Fig. 4B and Fig. S11), presumably because the OmpC-Mla system may have higher specificity for PG (Thong *et al.*, 2016). Since PE is the predominant PL species in the OM (Fig. S10) (Cronan, 2003), overexpressing the OmpC-Mla system may not effectively reduce the overall build-up of PLs caused by the loss of Tol-Pal function. Further to validating the putative PL transport function of the OmpC-Mla system, our observation here lends strong support to the notion that the Tol-Pal complex may be a major system for retrograde PL transport.

## Discussion

Our work reveals that the Tol-Pal complex plays an important role in maintaining OM lipid homeostasis, possibly via retrograde PL transport. Removing the system causes accumulation of excess PLs (over LPS) in the OM (Fig. 2). While pathways for anterograde PL transport remain to be discovered, this result indicates that PL flux to the OM may be intrinsically higher than that of LPS. Evidently, the ability to transport high levels of PLs to the OM allows cells to compensate for the loss of OMPs due to defects in assembly (Fig. 2). Our data suggest that cells maintain an excess flux of PLs to the OM in order to offset changes in the unidirectional assembly pathways for other OM components, and then return the PL surplus to the IM via retrograde transport. Having bidirectional PL transport therefore provides a mechanism to regulate and ensure the formation of a stable OM.

It is not clear whether the Tol-Pal complex directly mediates retrograde PL transport. It is formally possible that the effects we have observed on retrograde PL transport are due to indirect effects of removing the Tol-Pal complex on other OM processes. However, we have already shown that removing this complex does not affect the assembly of both OMPs and LPS, two major components in the OM (Fig. S2). Consistently, we have demonstrated that strains with impaired OMP assembly do not have defects in retrograde PL transport (Fig. 4A). We have also examined our strains under conditions where *tol-pal* mutants do not exhibit apparent division defects (Gerding *et al*., 2007); it is thus unlikely that there could be indirect effects on retrograde PL transport arising from the role of the Tol-Pal complex during cell division. Therefore, we believe that the Tol-Pal complex may directly mediate PL transport. One possibility is that this machine directly binds and transports lipids, even though there are no obvious lipid binding motifs or cavities found in available structures of the periplasmic components (Deprez C *et al*., 2005; Carr *et al*., 2000). The Tol-Pal complex is related to the ExbBD-TonB (Cascales *et al*., 2001; Celia *et al*., 2016), Agl-Glt (Faure *et al*., 2016), and Mot (Cascales *et al*., 2001; Thormann and Paulick, 2010) systems, each of which uses pmf-energized conformational changes to generate force for the uptake of metal-siderophores, for gliding motility, or to power flagella rotation, respectively. In addition, both the Tol-Pal and ExbBD-TonB complexes are hijacked by toxins (such as colicins) and bacteriophages to penetrate the OM (Cascales *et al*., 2007). It is therefore also possible that the Tol-Pal complex acts simply as a force generator to transport other PL-binding proteins across the periplasm, or perhaps bring the OM close enough to the IM for PL transfer to occur via hemifusion events. For the latter scenario, one can envision energized TolA pulling the OM inwards via its interaction with Pal, which is anchored to the inner leaflet of the OM (Godlewska *et al*., 2009). While it remains controversial, the formation of such “zones of adhesion”, or membrane contact sites, has previously been proposed (Bayer, 1991), and in fact, was suggested to be a mechanism for retrograde transport of native and foreign lipids (Jones and Osborn, 1977).

That the Tol-Pal complex is involved in retrograde PL transport also has significant implications for Gram-negative bacterial cell division. As part of the divisome, this system is important for proper OM invagination during septum constriction (Gerding *et al*., 2007; Yeh *et al*., 2010; Jacquier *et al*., 2015). How OM invagination occurs is unclear. Apart from physically tethering the IM and the OM, we propose that removal of PLs from the inner leaflet of the OM, possibly by the Tol-Pal complex, serves to locally reduce the surface area of the inner leaflet relative to the outer leaflet (McMahon and Gallop, 2005). According to the bilayer-couple model (Sheetz and Singer, 1974), this may then induce the requisite negative curvature in the OM at the constriction site, thus promoting formation of the new cell poles.

Given the importance of the Tol-Pal complex in OM stability and bacterial cell division, it would be an attractive target for small molecule inhibition. This is especially so in some organisms, including the opportunistic human pathogen *Pseudomonas aeruginosa*, where the complex is essential for growth (Dennis *et al*., 1996; Lo Sciuto *et al*., 2014). The lack of understanding of the true function of the Tol-Pal complex, however, has impeded progress. We believe that our work in elucidating a physiological role of this complex will accelerate efforts in this direction, and contribute towards the development of new antibiotics in our ongoing fight against recalcitrant Gram-negative infections.

### Experimental Procedures

#### Bacterial strains and growth conditions

All the strains used in this study are listed in Table S1. *Escherichia coli* strain MC4100 [*F*^*-*^*araD139* Δ*(argF-lac) U169 rpsL150 relA1 flbB5301 ptsF25 deoC1 ptsF25 thi*] (Casadaban, 1976) was used as the wild-type (WT) strain for most of the experiments. To achieve accumulation of phosphatidylserine (PS) in cells, a temperature-sensitive phosphatidylserine decarboxylase mutant (*psd2*), which accumulates PS at the non-permissive temperature, was used (Hawrot and Kennedy, 1978). NR754, an *araD*^+^ revertant of MC4100 (Ruiz et al., 2008), was used as the WT strain for experiments involving overexpression of *lpxC* from the arabinose-inducible promoter (P_BAD_). Δ*tolQ*, Δ*tolA* and Δ*tol-pal* deletions were constructed using recombineering (Datsenko and Wanner, 2000) and all other gene deletion strains were obtained from the Keio collection (Baba et al., 2006). Whenever needed, the antibiotic resistance cassettes were flipped out as described (Datsenko and Wanner, 2000). Gene deletion cassettes were transduced into relevant genetic background strains via P1 transduction (Silhavy *et al*., 1984). Luria-Bertani (LB) broth (1% tryptone and 0.5% yeast extract, supplemented with 1% NaCl) and agar were prepared as previously described (Silhavy *et al*., 1984). Strains were grown in LB medium with shaking at 220 rpm at either 30°C, 37°C, or 42°C, as indicated. When appropriate, kanamycin (Kan; 25 μg ml^-1^), chloramphenicol (Cam; 30 μg ml^-1^) and ampicillin (Amp; 125 μg ml^-1^) were added.

#### Plasmid construction

All the plasmids used in this study are listed in Table S2. Desired genes were amplified from MC4100 chromosomal DNA using the indicated primers (sequences in Table S3). Amplified products were digested with indicated restriction enzymes (New England Biolabs), which were also used to digest the carrying vector. After ligation, recombinant plasmids were transformed into competent NovaBlue (Novagen) cells and selected on LB plates containing appropriate antibiotics. DNA sequencing (Axil Scientific, Singapore) was used to verify the sequence of the cloned gene.

To generate *tolR*_*D23R*_ mutant construct, site-directed mutagenesis was conducted using relevant primers listed in Table S3 with pET23/42*tolR* as the initial template. Briefly, the entire template was amplified by PCR and the resulting PCR product mixture digested with DpnI for > 1 h at 37°C. Competent NovaBlue cells were transformed with 1 μl of the digested PCR product and plated onto LB plates containing ampicillin. DNA sequencing (Axil Scientific, Singapore) was used to verify the introduction of the desired mutation.

#### Analysis of [^32^P]-labelled lipid A

Mild acid hydrolysis was used to isolate lipid A as previously described (Zhou *et al*., 1999) with some modifications. 5-ml cultures were grown in LB broth (inoculated from an overnight culture at 1:100 dilution) containing [^32^P]-disodium phosphate (final 1 μCi ml^-1^; Perkin Elmer product no. NEX011001MC) till mid-log phase (OD600 ∼0.5 - 0.7). One MC4100 WT culture labelled with [^32^P] was treated with EDTA (25 mM pH 8.0) for 10 min prior to harvesting. Cells were harvested at 4,700 x *g* for 10 min, washed twice with 1 ml PBS (137 mM NaCl, 2.7 mM KCl, 10 mM Na_2_HPO_4_, 1.8 mM KH_2_PO_4_, pH 7.4) and suspended in PBS (0.32 ml) again. Chloroform (0.4 ml) and methanol (0.8 ml) were added and the mixtures were incubated at room temperature for 20 min with slow shaking (60 rpm) to make the one-phase Bligh-Dyer mixture (chloroform:methanol:water = 1:2:0.8). Mixtures were then centrifuged at 21,000 x *g* for 30 min. Pellets obtained were washed once with fresh one-phase Bligh-Dyer system (1 ml) and centrifuged as above. Resulting pellets were suspended in 0.45 ml of sodium acetate (12.5 mM, pH 4.5) containing SDS (1 %) and heated at 100°C for 30 min. After cooling to room temperature, chloroform and methanol (0.5 ml each) were added to create a two-phase Bligh-Dyer mixture (chloroform:methanol:water = 2:2:1.8). The lower (organic) phase of each mixture was collected after phase partitioning via centrifugation at 21,000 x *g* for 30 min. This was washed once with upper phase (0.5 ml) of freshly prepared two-phase Bligh-Dyer mixture and centrifuged as above. Finally, all the collected lower phases containing [^32^P]-labelled lipid A were air-dried overnight. Dried radiolabelled lipid A samples were suspended in 50 μl of chloroform:methanol (2:1) and equal amounts (∼1,000 cpm) of radioactivity were spotted on silica-gel coated TLC (Thin Layer Chromatography) plates (Merck). TLCs were developed in chambers pre-equilibrated overnight with solvent system chloroform:pyridine:98 % formic acid:water (50:50:14.6:5). TLC plates were air-dried overnight and later visualized by phosphor imaging (STORM, GE healthcare). The densitometric analysis of the spots obtained on the phosphor images of TLCs was carried out using ImageQuant TL analysis software (version 7.0, GE Healthcare). Average levels of hepta-acylated lipid A (expressed as a percentage of total lipid A in each sample) were obtained from three independent experiments.

#### Sucrose density gradient fractionation

Sucrose density gradient centrifugation was performed as previously described (Chng *et al*., 2010) with some modifications. For each strain, a 10/50-ml culture (inoculated from an overnight culture at 1:100 dilution) was grown in LB broth until OD_600_ reached ∼0.5 – 0.7. For radiolabeling, indicated radioisotopes were added from the start of inoculation. Cells were harvested by centrifugation at 4,700 x *g* for 10 min, suspended to wash once in 5 ml of cold Buffer A (Tris-HCl, 10 mM pH 8.0), and centrifuged as above. Cells were resuspended in 6 ml of Buffer B (Tris-HCl, 10 mM pH 8.0 containing 20% sucrose (w/w), 1 mM PMSF and 50 μg ml ^-1^ DNase I), and lysed by a single passage through a high pressure French press (French Press G M, Glen Mills) homogenizer at 8,000 psi. Under these conditions, lipid mixing between inner and outer membranes is minimal (Chng *et al*., 2010). Unbroken cells were removed by centrifugation at 4,700 x *g* for 10 min. The cell lysate was collected, and 5.5 ml of cell lysate was layered on top of a two-step sucrose gradient consisting of 40% sucrose solution (5 ml) layered on top of 65% sucrose solution (1.5 ml) at the bottom of the tube. All sucrose (w/w) solutions were prepared in Buffer A. Samples were centrifuged at 39,000 rpm for 16 h in a Beckman SW41 rotor in an ultracentrifuge (Model XL-90, Beckman). 0.8-ml fractions (usually 15 fractions) were manually collected from the top of each tube.

#### Analysis of OMP and LPS levels in isolated OMs

OM fragments were isolated from 50 ml of cells following growth, cell lysis and application of sucrose density gradient fractionation, as described above. Instead of manual fractionation, OM fragments (∼1 ml) were isolated from the 40%/65% sucrose solution interface by puncturing the side of the tube with a syringe. Buffer A (1 ml) was added to the OM fragments to lower the sucrose concentration and reduce viscosity. The OM fragments were then pelleted in a microcentrifuge at 21,000 x *g* for 30 min and then resuspended in 200 - 250 μl Buffer A. Protein concentrations of these OM preparations were determined using Bio-Rad *D*_*C*_ protein assay. The same amount of OM (based on protein content) for each strain was analyzed by reducing SDS-PAGE and immunoblotted using antibodies directed against OmpC, OmpF, LamB, BamA, LptE and LPS. For LPS quantification, five-fold serial dilutions of WT OMs were ran alongside the other OM samples as standards. Densitometric analysis of the LPS bands was carried out using ImageJ analysis software, and calibrated using ratio standard curves generated from the serial dilution standards (Pitre *et al*., 2007). LPS levels found in the OMs of indicated strains were normalized to WT. This quantification was performed three times for the same samples, and the average data was plotted.

#### Analysis of steady-state [^3^H]-glycerol-labelled PL distribution in IMs and OMs

To specifically label cellular PLs, 10-ml cells were grown at 37°C in LB broth (inoculated from an overnight culture at 1:100 dilution) containing [2-^3^H]-glycerol (final 1 μCi ml^-1^; Perkin Elmer product no. NET022L001MC) until OD_600_ reached ∼0.5 - 0.7. Once the desired OD_600_ was achieved, cultures were immediately mixed with ice-cold Buffer A containing CCCP (50 μM) to stop the labeling of the cultures. Cells were pelleted, lysed, and fractionated on sucrose density gradients, as described above. 0.8-ml fractions were collected from each tube, as described above, and 300 μl from each fraction was mixed with 2 ml of Ultima Gold scintillation fluid (Perkin Elmer, Singapore). Radioactivity ([^3^H]-count) was measured on a scintillation counter (MicroBeta^2®^, Perkin-Elmer). Based on [^3^H]-profiles, IM and OM peaks were identified and peak areas determined after background subtraction (average count of first 5 fractions was taken as background). For each strain, relative [^3^H]-PL levels in the IM and OM were expressed as a percentage of the sum in both membranes (see Fig. 2A upper panel). The average percent [^3^H]-PL in the OM for each strain (obtained from three independent experiments) was then compared to that for the WT strain to calculate fold changes (see Fig. 2B lower panel).

#### Determination of PL/LPS ratios in [^14^C]-acetate labelled OMs (see Fig. S5 for workflow and results)

To specifically label all cellular lipids (including LPS), 10-ml cells were grown at 37°C in LB broth (inoculated from an overnight culture at 1:100 dilution) containing [1-^14^C]-acetate (final 0.2 μCi ml^-1^; Perkin Elmer product no. NEC084A001MC) until OD_600_ reached ∼0.5 – 0.7. At this OD, cultures were transferred immediately to ice-cold Buffer A (5 ml), pelleted, lysed, and fractionated on sucrose density gradients, as described above. 0.8-ml fractions were collected from each tube, as described above, and 50 μl from each fraction was mixed with 2 ml of Ultima Gold scintillation fluid (Perkin Elmer, Singapore). Based on [^14^C]-profiles, IM and OM peaks were identified. OM fractions were then pooled, and treated as outlined below to differentially extract PLs and LPS for relative quantification within each OM pool. For each strain, the whole experiment was conducted and the OM PL/LPS ratio obtained three times.

Each OM pool (0.32 ml) was mixed with chloroform (0.4 ml) and methanol (0.8 ml) to make a one-phase Bligh-Dyer mixture (chloroform:methanol:water = 1:2:0.8). The mixtures were vortexed for 2 min and later incubated at room temperature for 20 min with slow shaking at 60 rpm. After centrifugation at 21,000 x *g* for 30 min, the supernatants (S1) were collected. The resulting pellets (P1) were washed once with fresh 0.95 ml one-phase Bligh-Dyer solution and centrifuged as above. The insoluble pellets (P2) were air dried and used for LPS quantification (see below). The supernatants obtained in this step (S2) were combined with S1 to get the combined supernatants (S3), which contained radiolabelled PLs. To these, chloroform (0.65 ml) and methanol (0.65 ml) were added to convert them to two-phase Bligh-Dyer mixtures (chloroform:methanol:water = 2:2:1.8). After a brief vortexing step, the mixtures were centrifuged at 3000 x *g* for 10 min to separate the immiscible phases, and the lower organic phases were collected. These were washed once with equal volumes of water and centrifuged as above, and the lower organic phases (containing radiolabelled PLs) recollected and air dried. Finally, the dried PLs were dissolved in 50 μl of a mixture of chloroform:methanol (2:1). Equal volumes (20 μl) of PL solutions were mixed with 2 ml of Ultima Gold scintillation fluid (Perkin Elmer, Singapore). The [^14^C]-counts were measured using scintillation counting (MicroBeta^2®^, Perkin-Elmer) and taken as the levels of PLs isolated from the OMs.

To quantify LPS, the P2 pellets were suspended in 2X reducing SDS-PAGE loading buffer (40 μl) and boiled for 10 min. Equal volumes (15 μl) were loaded and subjected to SDS PAGE (15% Tris.HCl). Gels were air-dried between porous films (Invitrogen) and exposed to the same phosphor screen along with standards (GE healthcare). To generate a standard curve for LPS quantification, the WT OM pellet sample was serially diluted two-fold and equal volumes of diluted samples were resolved on SDS-PAGE and dried as above. The densitometric analysis of bands (i.e. LPS from each OM) was carried out using ImageQuant TL analysis software (version 7.0, GE Healthcare). To allow proper comparison and quantification, the LPS gels from triplicate experiments were exposed on the same phosphor screen along with the standards (see Fig. S5).

For each strain, the arbitrary PL/LPS ratio in the OM was obtained by taking the levels of PLs (represented by [^14^C]-counts of PL fraction) divided by the LPS levels (represented by gel band density), averaged across three independent replicates (see Fig. 5C and Fig. 2B upper panel). The average PL/LPS ratio in the OM for each strain was then compared to that for the WT strain to calculate fold changes (see Fig. 2B lower panel).

#### Quantification of OM vesiculation

For each strain, 10-ml cells were grown at 37°C in LB broth (inoculated from an overnight culture at 1:100 dilution) containing [1-^14^C]-acetate (final 0.2 μCi ml^-1^; Perkin Elmer product no. NEC084A001MC) until OD_600_ reached ∼0.7. At this OD, cultures were harvested to obtain the cell pellets, and supernatants containing OM vesicles. Cell pellets were washed twice with Buffer A and finally suspended in the same buffer (0.2 ml). To obtain OM vesicles, supernatants were filtered through 0.45 μm filters followed by ultracentrifugation in a SW41.Ti rotor at 39,000 rpm for 1 h. Finally, the OM vesicles in the resulting pellets were washed and resuspended in 0.2 ml of Buffer A. Radioactive counts in cell pellets and OM vesicles were measured after mixing with 2 ml of Ultima Gold scintillation fluid (Perkin Elmer, Singapore). Radioactivity ([^14^C]-count) was measured on a scintillation counter (MicroBeta^2®^, Perkin-Elmer).

#### PG/CL turnover assay (pulse-chase and single time-point (2-h) analysis)

PG/CL turnover pulse-chase experiments were performed using the *psd2* background, which accumulated PS and PG/CL during growth at restrictive temperature. For each strain, cells were grown in 70 ml LB broth (inoculated from an overnight culture at 1:100 dilution) at the permissive temperature (30°C) until OD_600_ reached ∼0.15 - 0.2. The culture was then shifted for 4 h at the restrictive temperature (42°C) and labelled with [^32^P]-disodium phosphate (final 1 μCi ml^-1^) during the last 30 min at the restrictive temperature (42°C). After labeling, cells were harvested by centrifugation at 4,700 x *g* for 10 min, washed once with cold LB broth (10 ml) and centrifuged again at 4,700 x *g* for 10 min. Cells were then resuspended in fresh LB broth (70 ml) and the chase was started in the presence of non-radioactive disodium phosphate (1000-fold molar excess) at either the permissive temperature, with or without addition of carbonyl cyanide *m*-chlorophenyl hydrazone (CCCP; 50 μM), or at the restrictive temperature in the presence of hydroxylamine (HA; 10 mM). At the start (0 min) and different times (15, 30, 45, 90 and 120 min) during the chase, a portion of the culture (either 15 ml or 10 ml) was collected and mixed immediately with equal volume of ice-cold Buffer A containing CCCP (50 μM) and hydroxylamine (10 mM). Cells were harvested by centrifugation at 4,700 x *g* for 10 min and then resuspended in 6 ml of Buffer B containing CCCP (50 μM) and hydroxylamine (10 mM). Cells were lysed, and fractionated on sucrose density gradients, as described above. 0.8-ml fractions were collected from each tube, as described above. Fractions 7-9 and 12-14 contained the IM and OM fractions, respectively. To extract PLs from the IM and OM pools (2.4 ml), methanol (6 ml) and chloroform (3 ml) were added to make one-phase Bligh-Dyer mixtures. These were incubated at room temperature for 60 min with intermittent vortexing. Chloroform (3 ml) and sterile water (3 ml) were then added to generate two-phase Bligh-Dyer mixtures. After brief vortexing, the lower organic phases were separated from the top aqueous phases by centrifugation at 3,000 x *g* for 10 min. These were washed once with equal volumes of water and centrifuged as above, and the lower organic phases (containing radiolabelled PLs) recollected and air dried. Finally, the dried PLs were dissolved in 40 μl of a mixture of chloroform:methanol (2:1) and spotted onto silica-gel coated TLC plates (Merck). Equal amounts (in cpm) of radioactivity were spotted for each sample. TLCs were developed in pre-equilibrated chambers containing solvent system chloroform:methanol:water (65:25:4). TLC plates were dried, and visualized by phosphor imaging (STORM, GE healthcare). Densitometric analysis of the PL spots on the phosphor image of TLCs was conducted using the ImageQuant TL analysis software (version 7.0, GE Healthcare). The levels of each major PL species were expressed as a percentage of all detected PL species (essentially the whole lane), and plotted against time (see Fig. 3 and Fig. S8).

For single time-point analysis, 30-ml cultures were grown and labelled with [^32^P]-disodium phosphate (final 1 μCi ml^-1^) at the restrictive temperature. For strains harboring plasmids used for overexpressing OmpC-Mla components, arabinose (0.2 %) was added during growth at the permissive as well as restrictive temperatures. After washing and resuspension in fresh LB broth (30 ml), the chase was started in the presence of non-radioactive disodium phosphate (1000-fold molar excess) at the permissive temperature. At start (0 h) and 2 h during the chase, a portion of the culture (15 and 10 ml) was collected and processed similarly as pulse chase analysis described above. The levels of PG/CL in the membranes at each time point were expressed as a percentage of the sum of PE, PS and PG/CL. For each strain, IM and OM PG/CL turnover were expressed as the difference between percentage PG/CL levels at 0-h and 2-h time points divided by that at 0-h. Average PG/CL turnover values were obtained from three independent experiments conducted (see Fig. 4 and Fig. S9).

#### OM permeability assay

OM sensitivity against SDS/EDTA was judged by colony-forming unit (cfu) analyses on LB agar plates containing indicated concentrations of SDS/EDTA. Briefly, 5-ml cultures were grown (inoculated with overnight cultures at 1:100 dilution) in LB broth at 37°C until OD_600_ reached ∼1.0. Cells were normalized according to OD_600_, first diluted to OD_600_ = 0.1 (∼10^8^ cells), and then serial diluted in LB with seven 10-fold dilutions using 96-well microtiter plates (Corning). Two microliters of the diluted cultures were manually spotted onto the plates and incubated overnight at 37°C.

#### LpxC overexpression (growth curves and viability assay)

For each strain, a 10-ml culture was inoculated in LB broth supplemented with arabinose (0.2 %) from the overnight culture to make the initial OD_600_ of 0.05. Cells were grown at 37°C and the OD_600_ of the cultures was measured hourly. At the start of growth (0 h) and at 4 and 7 h during growth, 100 μl of cells were collected and then serial diluted in LB/cam with six 10-fold dilutions using 96-well microtiter plates (Corning). Five microliters of the non-diluted and diluted cultures were manually spotted on LB/cam agar plates (no arabinose). Plates were incubated overnight at 37°C.

#### IM (NADH activity) and OM marker (LPS) analysis during sucrose gradient fractionation

The inner membrane enzyme, NADH oxidase, was used as a marker for the IM; its activity was measured as previously described (Chng *et al*., 2010). Briefly, 30 μl of each fraction from the sucrose density gradient was diluted 4-fold with 20 mM Tris.HCl, pH 8.0 in a 96-well format and 120 μl of 100 mM Tris.HCl, pH 8.0 containing 0.64 mM NADH (Sigma) and 0.4 mM dithiothreitol (DTT, Sigma) was added. Changes in fluorescence over time due to changes in NADH (λex = 340 nm, λem = 465 nm) concentration was monitored using a plate reader (Perkin Elmer). The activity of NADH oxidase in pooled IM and OM fractions relative to the sum of these fractions was determined.

LPS was used as a marker for the OM and detected using LPS dot blots. OM fractions were pooled together and 2 μl of the fractions were spotted on nitrocellulose membranes (Bio-Rad). Spotted membranes were allowed to dry at room temperature for 1 h and then the membranes were probed with antibodies against LPS.

#### SDS-PAGE and immunoblotting

All samples subjected to SDS-PAGE were mixed with 2X Laemmli reducing buffer and boiled for 10 min at 100°C. Equal volumes of the samples were loaded onto the gels. Unless otherwise stated, SDS-PAGE was performed according to Laemmli using the 12% or 15% Tris.HCl gels (Laemmli, 1970). Immunoblotting was performed by transferring protein bands from the gels onto polyvinylidene fluoride (PVDF) membranes (Immun-Blot® 0.2 μm, Bio-Rad) using the semi-dry electroblotting system (Trans-Blot® TurboTM Transfer System, Bio-Rad). Membranes were blocked using 1X casein blocking buffer (Sigma). Mouse monoclonal α-OmpC antibody was a gift from Swaine Chen and used at a dilution of 1:5,000 (Khetrapal et al., 2015). Rabbit α-LptE (from Daniel Kahne) (Chng et al., 2010) and α-OmpF antisera (Rajeev Misra) (Charlson et al., 2006) were used at 1:5,000 dilutions. Rabbit α-BamA antisera (from Daniel Kahne) was used at 1:40,000 dilution. Rabbit α-LpxC antisera (generous gift from Franz Narberhaus) was used at 1:5,000 dilution. Mouse monoclonal α-LPS antibody (against LPS-core) was purchased from Hycult biotechnology and used at 1:5,000 dilutions. Rabbit polyclonal α-LamB antibodies was purchased from Bioss (USA) and used at 1:1,000 dilution. α-mouse IgG secondary antibody conjugated to HRP (from sheep) and α-rabbit IgG secondary antibody conjugated to HRP (from donkey) were purchased from GE Healthcare and used at 1:5,000 dilutions. Luminata Forte Western HRP Substrate (Merck Milipore) was used to develop the membranes and chemiluminescent signals were visualized by G:BOX Chemi XT 4 (Genesys version1.3.4.0, Syngene).

## Acknowledgments

We thank Zhi-Soon Chong for constructing the Δ*mlaC* allele, and Chee-Geng Chia for performing preliminary experiments. We are grateful to Swaine Chen (NUS), Rajeev Misra (Arizona State U), Daniel Kahne (Harvard U) and Franz Narberhous (RUHR Universitat Bochum) for their generous gifts of α-OmpC, α-OmpF, α-LptE and α-BamA, and α-LpxC antibodies, respectively. Finally, we thank William F. Burkholder (Institute of Molecular and Cell Biology) and Jean-Francois Collet (U Catholique de Louvain) for critical comments and suggestions on the manuscript. This work was supported by the National University of Singapore Start-up funding, the Singapore Ministry of Education Academic Research Fund Tier 1 and Tier 2 (MOE2013-T2-1-148) grants, and the Singapore Ministry of Health National Medical Research Council under its Cooperative Basic Research Grant (NMRC/CBRG/0072/2014) (all to S.-S.C.).

## Author contributions

R.S. performed all experiments described in this work; X.E.J. performed experiments related to LpxC overexpression; R.S. and S.-S.C. analyzed and discussed data; R.S. and S.-S.C. wrote the paper.

## References

Audet, A., Cole, R., and Proulx, P. (1975) Polyglycerophosphatide metabolism in *Escherichia coli*. Biochim Biophys Acta 380: 414–420.

Baba, T., Ara, T., Hasegawa, M., Takai, Y., Okumura, Y., Baba, M., et al. (2006) Construction of *Escherichia coli* K-12 in-frame, single-gene knockout mutants: the Keio collection. Mol Syst Biol 2: 2006.0008.

Bayer, M. E. (1991) Zones of membrane adhesion in the cryofixed envelope of *Escherichia coli*. J Struct Biol 107: 268–280.

Bernadac, A., Gavioli, M., Lazzaroni, J.C., Raina, S., Lloubes, R. (1998) *Escherichia coli tol-pal* mutants form outer membrane vesicles. J Bacteriol 180: 4872–4878.

Bernstein, A., Rolfe, B., and Onodera, K. (1972) Pleiotropic properties and genetic organization of the *tolA, B* locus of *Escherichia coli* K-12. J Bacteriol 112: 74–83.

Bishop, R.E. (2005) The lipid A palmitoyltransferase PagP: molecular mechanisms and role in bacterial pathogenesis. Mol Microbiol 57: 900–912.

Carr, S., Penfold, C.N., Bamford, V., James, R., and Hemmings, A.M. (2000) The structure of TolB, an essential component of the *tol*-dependent translocation system, and its protein-protein interaction with the translocation domain of colicin E9. Structure 8: 57–66.

Casadaban, M.J. (1976) Transposition and fusion of the *lac* genes to selected promoters in *Escherichia coli* using bacteriophage lambda and Mu. J Mol Biol 104: 541–555.

Cascales, E., Gavioli, M., Sturgis, J.N., and Lloubes, R. (2000) Proton motive force drives the interaction of the inner membrane TolA and outer membrane Pal proteins in *Escherichia coli*. Mol Microbiol 38: 904–915.

Cascales, E., Lloubes, R., and Sturgis, J.N. (2001) The TolQ-TolR proteins energize TolA and share homologies with the flagellar motor proteins MotA-MotB. Mol Microbiol 42: 795–807.

Cascales, E., Buchanan, S.K., Duche, D., Kleanthous, C., Lloubes, R., Postle, K., et al. (2007) Colicin biology. Microbiol Mol Biol Rev 71: 158–229.

Celia, H., Noinaj, N., Zakharov, S.D., Bordignon, E., Botos, I., Santamaria, M., et al. (2016) Structural insight into the role of the Ton complex in energy transduction. Nature 538: 60–65.

Charlson, E.S., Werner, J.N., and Misra, R. (2006) Differential effects of *yfgL* mutation on *Escherichia coli* outer membrane proteins and lipopolysaccharide. J Bacteriol 188: 7186–7194.

Chong, Z.S., Woo, W.F., and Chng, S.S. (2015) Osmoporin OmpC forms a complex with MlaA to maintain outer membrane lipid asymmetry in *Escherichia coli*. Mol Microbiol 98: 1133–1146.

Chng, S.S., Gronenberg, L.S., and Kahne, D. (2010) Proteins required for lipopolysaccharide transport in *Escherichia coli* form a transenvelope complex. Biochemistry 49: 4565–4567.

Clavel, T., Lazzaroni, J.C., Vianney, A., and Portalier, R. (1996) Expression of the *tolQRA* genes of *Escherichia coli* K-12 is controlled by the RcsC sensor protein involved in capsule synthesis. Mol Microbiol 19: 19–25.

Cronan, J.E. (2003) Bacterial membrane lipids: where do we stand? Annu Rev Microbiol 57: 203–224.

Dalebroux, Z.D., Matamouros, S., Whittington, D., Bishop, R.E., and Miller, S.I. (2014) PhoPQ regulates acidic glycerophospholipid content of the *Salmonella typhimurium* outer membrane. Proc Natl Acad Sci USA 111: 1963–1968.

Datsenko, K.A., and Wanner, B.L. (2000) One-step inactivation of chromosomal genes in *Escherichia coli* K-12 using PCR products. Proc Natl Acad Sci USA 97: 6640–6645.

Dennis, J.J., Lafontaine, E.R., and Sokol, P.A. (1996) Identification and characterization of the *tolQRA* genes of *Pseudomonas aeruginosa*. J Bacteriol 178: 7059–7068.

Deprez, C., Lloubes, R., Gavioli, M., Marion, D., Guerlesquin, F., and Blanchard, L. (2005) Solution structure of the *E. coli* TolA C-termical domain reveals conformational changes upon binding to the phage g3p N-terminal domain. J Mol Biol 346: 1047–1057.

Donohue-Rolfe, A.M., and Schaechter, M. (1980) Translocation of phospholipids from the inner to the outer membrane of *Escherichia coli*. Proc Natl Acad Sci USA 77: 1867–1871.

Ekiert, D.C., Bhabha, G., Isom, G.L., Greenan, G., Ovchinnikov, S., Henderson, I.R., et al. (2017) Architectures of lipid transport systems for the bacterial outer membrane. Cell 169: 273–285.e17.

Faure, L.M., Fiche, J.B., Espinosa, L., Ducret, A., Anantharaman, V., Luciano, J., et al. (2016) The mechanism of force transmission at bacterial focal adhesion complexes. Nature 539: 530–535.

Fuhrer, F., Langklotz, S., and Narberhaus, F. (2006) The C-terminal end of LpxC is required for degradation by the FtsH protease. Mol Microbiol 59: 1025–1036.

Gerding, M.A., Ogata, Y., Pecora, N.D., Niki, H., and de Boer, P.A.J. (2007) The trans-envelope Tol-Pal complex is part of the cell division machinery and required for proper outer-membrane invagination during cell constriction in *E. coli*. Mol Microbiol 63: 1008–1025.

Germon, P., Ray, M.C., Vianney, A., and Lazzaroni, J.C. (2001) Energy-dependent conformational changes in the TolA protein of *Escherichia coli* involves its N-terminal domain, TolQ, and TolR. J Bacteriol 183: 4110–4114.

Gresock, M.G., Kastead, K.A., and Postle, K. (2015) From homodimer to heterodimer and back: elucidating the TonB energy transduction cycle. J Bacteriol 197: 3433–3445.

Godlewska, R., Wisniewska, K., Pietras, Z., and Jagusztyn-Krynicka, E.K. (2009) Peptidoglycan-associated lipoprotein (Pal) of Gram-negative bacteria: function, structure, role in pathogenesis and potential application in immunoprophylaxis. FEMS Microbiol Lett 298: 1–11.

Gully, D., and Bouveret, E. (2006) A protein network for phospholipid synthesis uncovered by a variant of the tandem affinity purification method in *Escherichia coli*. Proteomics 6: 282–293.

Guzman, L.M., Belin, D., Carson, M.J., and Beckwith, J. (1995) Tight regulation, modulation, and high-level expression by vectors containing the arabinose PBAD promoter. J Bacteriol 177: 4121–4130.

Hagan, C.L., Silhavy, T.J., and Kahne, D. (2011) β-barrel membrane protein assembly by the Bam complex. Annu Rev Biochem 80: 189–210.

Hawrot, E., and Kennedy, E.P. (1978) Phospholipid composition and membrane function in phosphatidylserine decarboxylase mutants of *Escherichia coli*. J Biol Chem 253: 8213–8220.

Hirschberg, C.B., and Kennedy, E.P. (1977) Mechanims of the enzymatic synthesis of cardiolipin in *Escherichia coli*. Proc Natl Acad Sci USA 69: 648–651.

Jacquier, N., Frandi, A., Viollier, P.H., and Greub, G. (2015) Disassembly of a medial transenvelope structure by antibiotics during intracellular division. Chem Biol 22: 1217–1227.

Jones, N.C., and Osborn, M.J. (1977) Translocation of phospholipids between the outer and inner membranes of *Salmonella typhimurium*. J Biol Chem 252: 7405–7412.

Kanemasa, Y., Akamatsu, Y., and Nojima, S. (1967) Composition and turnover of the phospholipids in *Escherichia coli*. Biochim Biophys Acta 144: 382–390.

Kanfer, J., and Kennedy, E.P. (1963) Metabolism and function of bacterial lipids I. Metabolism of phospholipids in *Escherichia coli* B. J Biol Chem 238: 2919–2922.

Khetrapal, V., Mehershahi, K., Rafee, S., Chen, S., Lim, C.L., and Chen, S.L. (2015) A set of powerful negative selection systems for unmodified Enterobacteriaceae. Nucleic Acids Res 43: e83.

Laemmli, U.K. (1970) Cleavage of structural proteins during the assembly of the head of bacteriophage T4. Nature 227: 680–685.

Langley, K.E., Hawrot, E., and Kennedy, E.P. (1982) Membrane assembly: movement of phosphatidylserine between the cytoplasmic and outer membranes of *Escherichia coli*. J Bacteriol 152: 1033–1041.

Lazzaroni, J.C., and Portalier, R.C. (1981) Genetic and biochemical characterization of periplasmic-leaky mutants of *Escherichia coli* K-12. J Bacteriol 145: 1351–1358.

Lloubes, R., Cascales, E., Walburger, A., Bouveret, E., Lazdunski, C., Bernadac, A., et al. (2001) The Tol-Pal proteins of the *Escherichia coli* cell envelope: an energized system required for outer membrane integrity? Res Microbiol 152: 523–529.

Lo Sciuto, A., Fernandez-Pinar, R., Bertuccini, L., Losi, F., Superti, F., Imperi, F. (2014) The periplasmic protein TolB as a potential drug target in *Pseudomonas aeruginosa*. PLoS One 9: e103784.

Malinverni, J.C., and Silhavy, T.J. (2009) An ABC transport system that maintains lipid asymmetry in the Gram-negative outer membrane. Proc Natl Acad Sci USA 106: 8009–8014.

McMahon, H.T., and Gallop, J.L. (2005) Membrane curvature and mechanisms of dynamic cell membrane remodelling. Nature 438: 590–596.

Nikaido, H. (2003) Molecular basis of bacterial outer membrane permeability revisited. Microbiol Mol Biol Rev 67: 593–656.

Nakayama, T., and Zhang-Akiyama, Q.M. (2016) *pqiABC* and *yebST*, putative mce operons of *Escherichia coli*, encode transport pathways and contribute to membrane integrity. J Bacteriol 199: e00606–16.

Okuda, S., Sherman, D.J., Silhavy, T.J., Ruiz, N., and Kahne, D. (2016) Lipopolysaccharide transport and assembly at the outer membrane: the PEZ model. Nat Rev Microbiol 14: 337–345.

Okuda, S., and Tokuda, H. (2011) Lipoprotein sorting in bacteria. Annu Rev Microbiol 65: 239–259.

Pitre, A., Pan, Y., Pruett, S., and Skalli, O. (2007) On the use of ratio standard curves to accurately quantitate relative changes in protein levels by western blot. Anal Biochem 361: 305–307.

Ruiz, N., Chng, S.S., Hinikera, A., Kahne, D., and Silhavy, T.J. (2010) Nonconsecutive disulphide bond formation in an essential integral outer membrane protein. Proc Natl Acad Sci USA 107: 12245–12250.

Ruiz, N., Falcone, B., Kahne, D., and Silhavy, T.J. (2005) Chemical conditionality: a genetic strategy to probe organelle assembly. Cell 121: 307–317.

Ruiz, N., Gronenberg, L.S., Kahne, D., and Silhavy, T.J. (2008) Identification of two inner-membrane proteins required for the transport of lipopolysaccharide to the outer membrane of *Escherichia coli*. Proc Natl Acad Sci USA 105: 5537–5542.

Satre, M., and Kennedy, E.P. (1978) Identification of bound pyruvate essential for the activity of phosphatidylserine decarboxylase of *Escherichia coli*. J Biol Chem 253: 479–483.

Schulman, H., and Kennedy, E.P. (1977) Relation of turnover of membrane phospholipids to synthesis of membrane-derived oligosaccharides of *Escherichia coli*. J Biol Chem 252: 4250–4255.

Sheetz, M.P., and Singer, S.J. (1974) Biological membranes as bilayer couples. A molecular mechanism of drug-erythrocyte interactions. Proc Natl Acad Sci USA 71: 4457–4461.

Silhavy, T.J., Berman, M.L., and Enquist, L.W. (1984) Experiments with Gene fusions (Cold Spring Harbor Laboratory Press, Cold Spring Harbor, New York).

Sturgis, J.N. (2001) Organisation and evolution of the *tol-pal* gene cluster. J Mol Microbiol Biotechnol 3: 113–122.

Thong, S., Ercan, B., Torta, F., Fong, Z.Y., Wong, H.Y., Wenk, M.R., et al. (2016) Defining key roles for auxillary proteins in an ABC transporter that maintains bacterial outer membrane lipid asymmetry. eLife 5: e19042.

Thormann, K.M., and Paulick, A. (2010) Tuning the flagellar motor. Microbiology 156: 1275–1283.

Vines, E.D., Marolda, C.L., Balachandran, A., and Valvano, M.A. (2005) Defective O-antigen polymerization in *tolA* and *pal* mutants of *Escherichia coli* in response to extracytoplasmic stress. J Bacteriol 187: 3359–3368.

Walburger, A., Lazdunski, C., and Corda, Y. (2002) The Tol/Pal system function requires an interaction between the C-terminal domain of TolA and the N-terminal domain of TolB. Mol Microbiol 44: 695–708.

Witty, M., Sanz, C., Shah, A., Grossmann, J.G., Mizuguchi, K., Perham, R.N., et al. (2002) Structure of the periplasmic domain of *Pseudomonas aeruginosa* TolA: evidence for an evolutionary relationship with the TonB transporter protein. EMBO J 21: 4207–4218.

Wu, T., Melinverni, J., Ruiz, N., Kim, S., Silhavy, T.J., and Kahne, D. (2005) Identification of a multicomponent complex required for outer membrane biogenesis in *Escherichia coli*. Cell 121: 235–245.

Wu, T., McCandlish, A.C., Gronenberg, L.S., Chng, S.S., Silhavy, T.J., and Kahne, D. (2006) Identification of a protein complex that assembles lipopolysaccharide in the outer membrane of *Escherichia coli*. Proc Natl Acad Sci USA 103: 11754–11759.

Yeh, Y.C., Comolli, L.R., Downing, K.H., Shapiro, L., and McAdams, H.H. (2010) The *Caulobacter* Tol-Pal complex is essential for outer membrane integrity and the positioning of a polar localization factor. J Bacteriol 192: 4847–4858.

Yem, D.W., and Wu, H.C. (1978) Physiological characterization of an *Escherichia coli* mutant altered in the structure of murein lipoprotein. J Bacteriol 133: 1419–1426.

Yokoto, K., and Kito, M. (1982) Transfer of the phosphatidyl moiety of phosphatidylglycerol to phosphatidylethanolamine in *Escherichia coli*. J Bacteriol 151: 952–961.

Zhou, Z., Lin, S., Cotter, R.J., and Raetz, C.R.H. (1999) Lipid A modifications characteristic of *Salmonella typhimurium* are induced by NH4VO3 in *Escherichia coli* K-12. J Biol Chem 274: 18503–18514.

